# The diel light cycle modulates microcystin content in continuous cultures and natural communities of *Microcystis*: Implications for bloom monitoring

**DOI:** 10.64898/2026.10.24.761440

**Authors:** Isabella J. Maggard, Robbie M. Martin, Kelly E. Joyce, Zac Triumph, Emily M. Garlock, Gregory L. Boyer, Justin D. Chaffin, David Talmy, Steven W. Wilhelm

## Abstract

Harmful algal blooms dominated by *Microcystis* spp. threaten fisheries, recreation, and drinking-water supplies because many strains produce the hepatotoxin microcystin. Monitoring programs compare microcystin concentrations among samples collected at different times of day, yet the extent that diel variation affects these estimates remains poorly resolved. We tested whether microcystin concentrations varied over a diel cycle under laboratory conditions and evaluated the effect of short-term sunlight exposure on a natural bloom community. Axenic *Microcystis aeruginosa* PCC 7806 was maintained in cyclostats with controlled nutrients and temperature under a 16 h light:8 h dark cycle. Microcystin concentrations normalized to culture volume (Μg L^-1^) and biovolume (ag μm^-3^) were significantly higher (*p* ≤ 0.05) at the end of the dark period (64 ± 8 Μg MC L^-1^; 630 ± 40 ag MC μm^-3^) than at the end of the light period (45 ± 7 Μg MC L^-1^; 390 ± 60 ag MC μm^-3^), and this pattern was reproducible across consecutive days. Cell abundance and photosynthetic efficiency remained constant, indicating variation was not explained by changes in growth or photosynthetic performance. In parallel, water from a Lake Erie *Microcystis* bloom was exposed to sunlight or paired dark controls, and microcystin concentrations declined ∼19% after just 60 min of sunlight exposure (p = 0.048). Together, these results demonstrate that microcystin measurements can vary predictably across the diel cycle and that light contributes to daytime declines. Although the underlying mechanism(s) remain unresolved, time of day should be considered, or standardized, when monitoring microcystins.

## MAIN NARRATIVE

Globally, harmful algal blooms (HABs) disrupt fisheries, aquaculture, and drinking water supplies. *Microcystis*, a cyanobacterium capable of producing hepatotoxic microcystins, dominates many of these blooms (Harke et al., 2016). For example, a *Microcystis*-dominated HAB in Lake Erie led to a temporary shutdown of Toledo’s (Ohio) water supply in 2014, affecting more than 400,000 residents and costing an estimated $65 million dollars (Bingham et al., 2015; Steffen et al., 2017). Such events highlight the importance of accurately characterizing bloom toxicity, which is mainly assessed by measuring microcystin concentrations. To this end, microcystins are routinely monitored in freshwater supplies globally (Franklin et al., 2026).

Microcystin monitoring programs are conducted by academic institutions, government agencies, and drinking water treatment plants. Monitoring programs involve sample collection from drinking water source intakes, beaches, and watercraft in the open water. Survey frequency can vary dramatically, ranging from daily to weekly or less often. Moreover, collection times may not be consistent from sample to sample and can be constrained by local weather patterns. Yet, collectively, microcystin measurements are often treated as representative of a population regardless of when samples were collected. This raises an important question: *does the time of sample collection influence measured microcystin concentration?*

Environmental variability complicates this question, as light, temperature, and nutrient availability can influence microcystin production (Wilhelm et al., 2020). Most laboratory studies of *Microcystis* rely on batch cultures, which are limited by changing growth conditions over time. Continuous cultures (chemostats) maintain constant environmental and chemical conditions, but many chemostat studies of *Microcystis* use continuous light, which does not reflect the natural diel (day/night) cycle (Oh et al., 2000). Cyclostats (Cosper, 1982; Rhee et al., 1981) address this limitation by maintaining chemically stable conditions and growth rates while incorporating a diel cycle. Here, we investigated diel variation of microcystin concentrations in continuous cultures of *Microcystis aeruginosa* PCC 7806 maintained under controlled temperature and nutrient conditions. We used cyclostats maintained under a 16 h light:8 h dark cycle at 50 μmol photons m^-2^ s^-1^ to measure changes in microcystin concentration at the end of the dark period (“dark”) and the end of the light period (“light”). We also investigated the effects of natural sunlight on a Lake Erie *Microcystis* bloom by measuring microcystin concentrations following short-term sunlight exposure.

Intracellular microcystin concentrations, normalized to cell biovolume (ag of microcystin per cell volume; ag MC μm^-3^), followed a repeatable diel pattern across the cyclostat experiment with an average cellular content of 390 ± 60 ag MC μm^-3^ after 16 hours of light and nearly doubled to 630 ± 40 ag MC μm^-3^ following 8 hours of darkness (Figure 1A, see supplementary methods for experimental details). The difference in ag MC μm^-3^ between dark and light was significant when data from all three days were pooled (Figure 1B; *p* = 0.026). Microcystin concentrations normalized to cell concentration (Μg MC cell^-1^) and total sample volume (Μg MC L^-1^) were also significantly different between light and dark (*p* ≤ 0.05; Supplemental Figure 1). Cell concentration and photosynthetic efficiency (*F*_*v*_*/F*_*m*_) did not change between the end of light and end of dark periods (Supplemental Figure 2A, 2B, 2E, and 2F), suggesting that the changes in microcystin concentration were not associated with changes in growth rate, cell abundance, or photosynthetic efficiency. Cell size increased significantly at the end of the light period (*p = 0*.*028*; Supplemental Figure 2C and 2D), supporting biovolume normalization to account for changes in cell size. Together, these results demonstrate consistent nighttime accumulation and daytime decline in measured microcystin concentrations. All data are presented in the supplemental tables.

**Figure 1:**
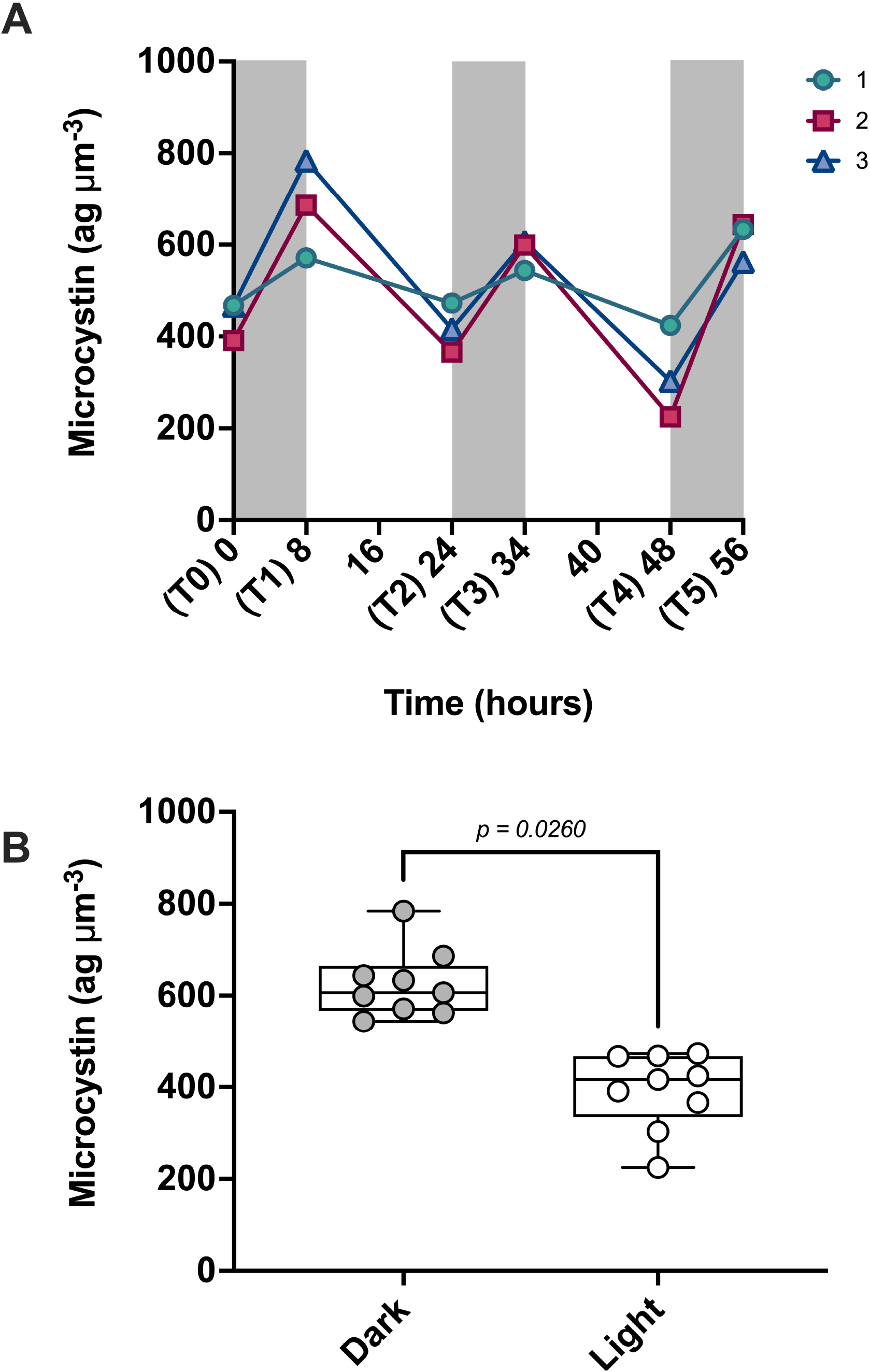
Diel variation in microcystin concentration in axenic continuous cultures of *M. aeruginosa* PCC 7806. (A) Microcystin concentration (normalized by biovolume) over three consecutive days. Gray shading indicates the dark period, and each color represent biological replicates. Differences among timepoints were evaluated using a repeated-measures ANOVA, with all pairwise timepoint comparisons significant (*p* ≤ 0.05). (B) Microcystin concentrations replotted by sampling period. Each dot represents a measurement from an individual cyclostat collected at the end of the dark (“dark”) or end of the light (“light”) periods on one of three consecutive days. Boxplots show the median and interquartile range. For statistical analysis, measurements from the three consecutive days were averaged within each cyclostat to yield a single value for each sampling period, with each cyclostat treated as an independent sample unit (n = 3). Differences between sampling periods were assessed using a paired t-test. Microcystin concentrations were significantly lower at the end of the light period (*p* = 0.026).

To test whether sunlight contributes to this pattern in natural *Microcystis*, we performed a short-term sunlight exposure experiment using an active *Microcystis* community collected from Lake Erie (Ohio, USA) during bloom conditions in August 2026. Briefly, homogenized lake water containing natural communities was distributed into 2-L polycarbonate bottles. Following overnight incubation, microcystin concentrations were measured before local sunrise (0 min sunlight exposure). Bottles were then wrapped in black plastic to prevent sunlight exposure until solar noon (∼1:00PM), when they were unwrapped and exposed to direct sunlight (∼1800 μmol photons m^-2^ s^-1^) for 30 or 60 min. Paired dark controls remained wrapped throughout the exposure period and were used to account for changes unrelated to light exposure.

The baseline microcystin concentration prior to the sunlight exposure was 2.5 ± 0.4 μg L^-1^. Measured microcystin concentrations decreased linearly with increasing duration of sunlight exposure (Figure 2B), decreasing to 2.2 ± 0.2 μg L^-1^ after 30 mins and 2.0 ± 0.2 μg L^-1^ after 60 mins, which was a significant difference (*p* = 0.048; Figure 2A). Microcystin concentrations in the paired dark controls (2.5 ± 0.1 μg L^-1^) did not differ significantly from the 0-min baseline (*p* = 0.101), indicating that the observed decrease was associated with sunlight exposure rather than duration time alone. The short-term response in natural lake water complements the diel pattern observed in lab cultures and suggests that light exposure is associated with changes in measured microcystin concentrations.

**Figure 2:**
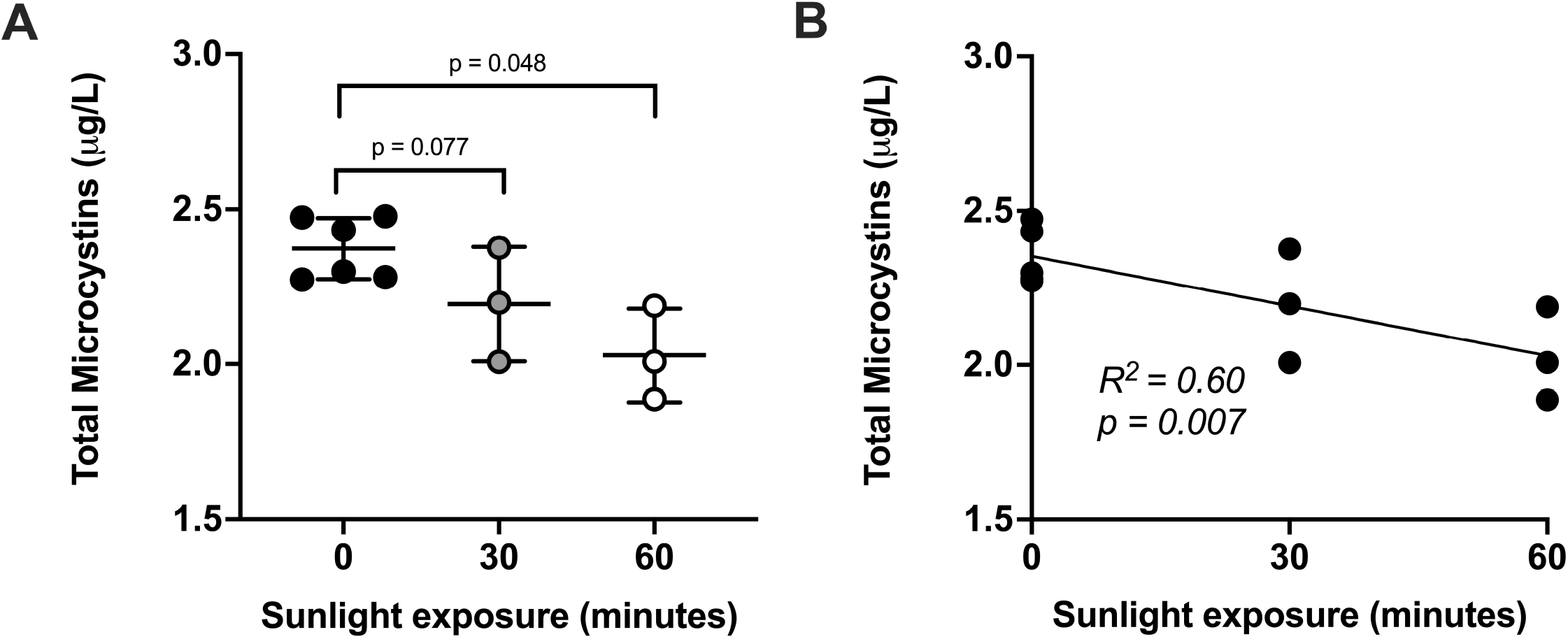
Short-term sunlight exposure is associated with changes in microcystin concentrations in natural communities. (A) Microcystin concentrations (normalized to total sample volume) measured before sunlight exposure (0 min) and following 30 or 60 min of direct sunlight exposure (∼1800 μmol photons m^-2^ s^-1^). Each point represents an independent biological replicate. Comparison between 0 and 30 min and 0 and 60 min were analyzed separately using a paired t-test (see supplementary methods for details about statistics). Microcystin concentrations decreased after 30 minutes of sunlight exposure, but the decrease was not significant (*p* = 0.077). Concentrations were significantly lower after 60 minutes (*p* = 0.048). (B) Simple linear regression of microcystin concentration against amount of sunlight exposure. Each point represents an independent biological replicate. The relationship was significant (*p* = 0.007) and explained 60% of the observed variation (*R*^*2*^ = 0.60).

Collectively, our observations demonstrate that measurable microcystin concentrations can vary throughout the diel cycle due to light exposure. The continuous culture experiment shows this pattern occurred when other environmental conditions were constrained, while the natural community experiment demonstrates that short-term sunlight exposure is associated with changes in microcystin concentration. These studies do not reveal the mechanism nor distinguish whether the observed changes reflect differences in microcystin production, release of microcystins to extracellular phase, degradation, or increased microcystin protein binding intracellularly resulting in non-detectable microcystins (Meissner et al., 2013; Zilliges et al., 2011). Rather, they demonstrate that the microcystins measured in a sample can depend on the time when that sample was collected. To this end, future microcystin monitoring programs should consider (and standardize when possible) the time of day when experimental or monitoring efforts are underway, as variability in microcystin concentrations may be associated with the duration of sunlight exposure. This could be particularly problematic when later day estimates are made that record low toxin concentrations relative to what may have existed just a few hours earlier.

## ACKNOWLEDGEMENTS

This work was supported by funds from the National Science Foundation (1840715, 2520197), the Allen Family Philanthropies (#202507-18146), and the National Institute of Environmental Health Sciences (1P01ES023-28 939-01). The authors thank Gary LeCleir for thoughts and comments on this project and Xiaocun Sun for advice on statistical analysis.

