## Supplementary figures and images for "The diel light cycle modulates microcystin content in continuous cultures and natural communities of *Microcystis*: Implications for bloom monitoring"

### supplemental figure 1

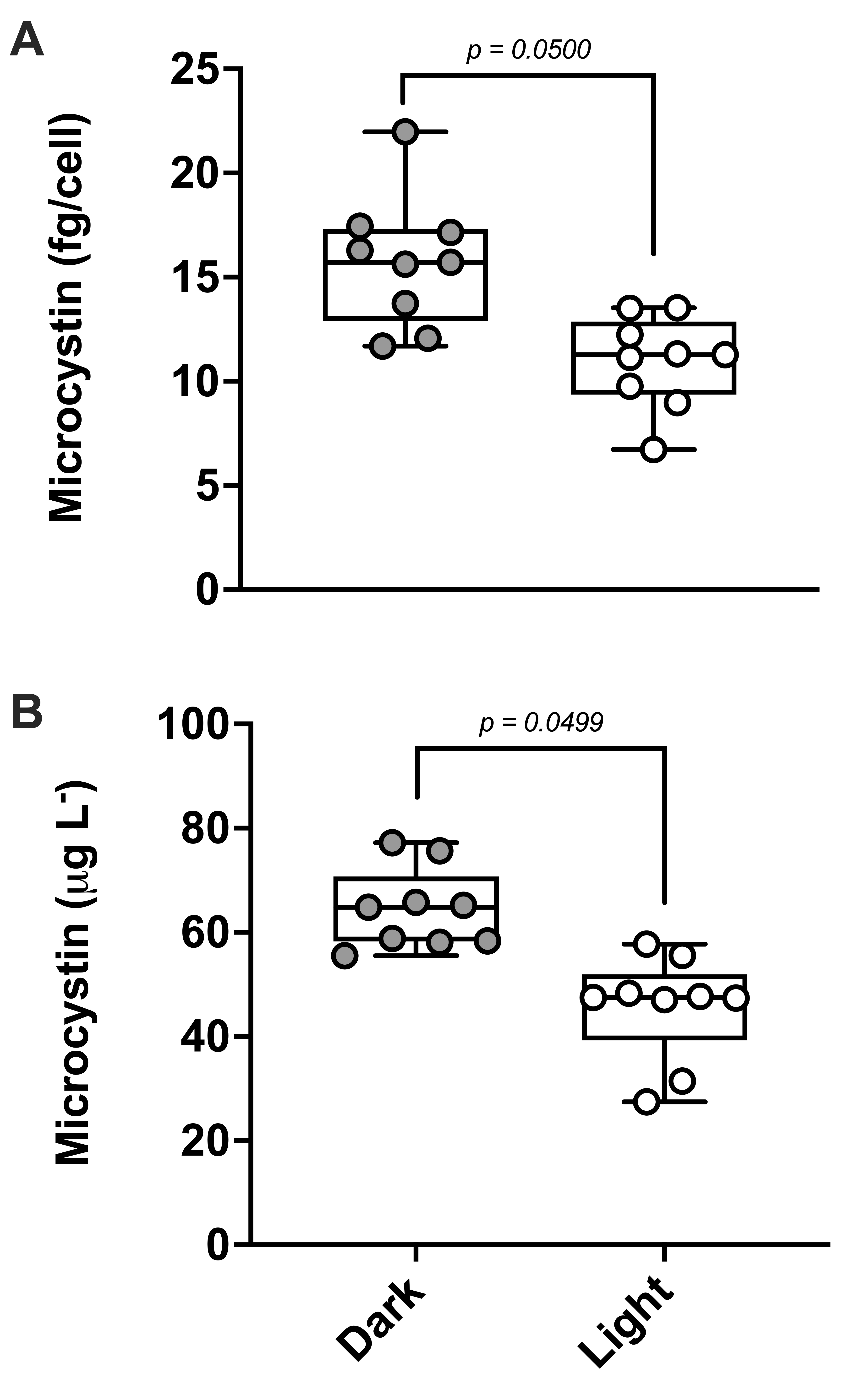

### supplemental figure 2A

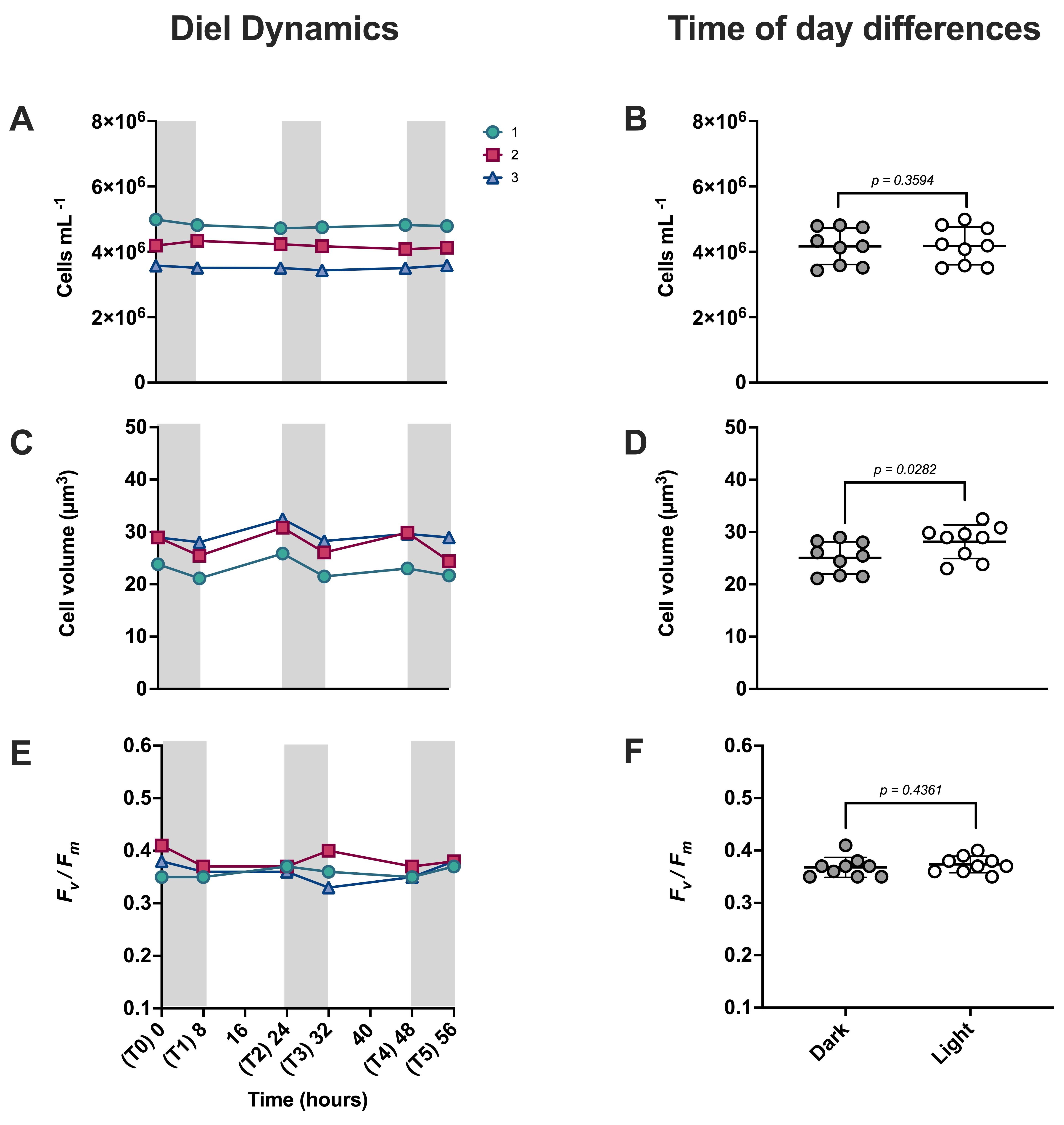
