## supplementary methods for "The diel light cycle modulates microcystin content in continuous cultures and natural communities of *Microcystis*: Implications for bloom monitoring"

**SUPPLEMENTAL FIGURE LEGENDS**

**Supplemental Figure 1: Microcystin concentrations normalized per cell (A) and per total sample volume in axenic continuous cultures of *M. aeruginosa* PCC 7806. (B).** Each dot represents a measurement from an individual cyclostat collected at the end of the dark (“dark”) or end of the light (“light”) periods on one of three consecutive days. Boxplots show the median and interquartile range. For statistical analysis, measurements from the three consecutive days were averaged within each cyclostat to yield a single value for each sampling period, with each cyclostat treated as an independent sample unit (n = 3). Differences between sampling periods were assessed using a paired t-test. Microcystin concentrations were significantly higher at the end of the dark period (*paired t-test, p ≤ 0.05*) for all normalization methods.

**Supplemental Figure 2: Diel dynamics in *Microcystis aeruginosa* PCC 7806 physiology in continuous cultures.** Diel dynamics of cell concentration (A), cell size (B), and photosynthetic efficiency (*F_v_/F_m_*; E). Gray shading indicates the dark period, and each color represent biological replicates. Data for cell concentration (B), cell size (D), and *F_v_/F_m_* (F) are replotted by sampling period. Each dot represents a measurement from an individual cyclostat collected at the end of the dark (“dark”) or end of the light (“light”) periods on one of three consecutive days. For statistical analysis, measurements from the three consecutive days were averaged within each cyclostat to yield a single value for each sampling period, with each cyclostat treated as an independent sample unit (n = 3). Differences between sampling periods were assessed using a paired t-test. Cell size increased at the end of the light period (*p* = 0.028).

**SUPPLEMENTAL METHODS**

*Continuous cultures.* To determine whether microcystin varied across the diel cycle under controlled conditions, we used continuous culture systems containing the toxigenic strain *M. aeruginosa* PCC 7806 modified to incorporate a day/night diel cycle (Martin et al., 2020). Cultures were grown in modified CT medium (Wilhelm, 2017) with a total N concentration of 161 µM (supplied as a mix of Ca(NO_3_)_2_ and KNO_3_) and an N:P ratio of 10. Three independent cyclostats were maintained in the same incubator at 26° C and a light intensity of 50 µmol photons m⁻² s⁻¹, with dilution rates of 0.25 d⁻¹ controlled by independent peristaltic pumps. Cultures were acclimated for approximately four weeks before sampling to allow steady-state conditions to be established. Samples were collected once cell concentrations remained stable (coefficient of variance for each individual cyclostat were all < 5% over a five-day period). Cyclostats were set on a 16:8 light:dark cycle, which represents early cyanobacterial bloom conditions for Lake Erie (day length is based on June 2, 2022, for Cleveland, OH). Cultures were sampled for three consecutive days at the end of the light period (we term as “light”) and at the end of the dark period (we term as “dark”).

*Cell concentration and size.* For each sampling event, cell concentration and mean cell diameter were determined. Cell concentration was measured using a Beckman Coulter CytoFLEX flow cytometer following the protocol published in *protocols.io* (Wren et al., 2026). Mean cell diameter was measured using a FlowCam 8000 with a 20× objective and FOV50 flowcell, with a minimum of approximately 15,000 cells analyzed per sample. Mean cell volume was calculated from the predicted mean diameter assuming spherical cells.

*Measurement of microcystin.* 50 mL of culture was collected at each time point, filtered onto 47-mm GF-75 glass fiber filters (Advantec) by vacuum filtration, immediately flash-frozen in liquid nitrogen, and stored at −80° C until analysis.

Quantification of microcystins for continuous culture experiments was performed via UPLC-MS/MS using a modified version of EPA method 544 (EPA, 2015). Eight common microcystins (RR, YR, dLR, LR, mLR, LA, LY, and LF) and nodularin-R were quantified using a Waters ACQUITY H-Class UPLC system (Waters Corporation, Milford, MA, USA) including a QSM solvent delivery system coupled to a FTN sample manager paired with a PDA UV-vis spectrometer and a TQD mass spectrometer that includes one quantitation ion and two confirmation ions for each analyte. An acidified (0.1% formic acid) 30-70% aqueous acetonitrile gradient at a flow rate of 0.2 mL min^-1^ was used with a 2.1 x 150 mm Ace-3 (3μm) C-18 column (MacMod Analytical, Chadds Ford, PA). UV-Vis response on the PDA was used for QAQC to ensure that the MS/MS response of the microcystin standards was acceptable. ESI with a capillary voltage of 3.5 kV, source temperature of 80°C, and desolvation gas temperature of 200°C was used with a flow of 550 L/hr of desolvation gas and 50 L/hr of cone gas. Additional method details including quantitation ions, fragmentation products, collision energies and cone voltages for each analyte can be seen in Supplemental Table 1. MCs were identified using the confirmation ion ratios and retention times as determined using standards (Supplemental Table 1). Detection limits varied between congeners and ranged from 0.18–2.73 (RR and LA respectively) µg microcystins L^-1^.

For the Lake Erie microcosm experiments, microcystins were measured using a modified version of the LC-MS/MS method from Birbeck et al. (2019). Microcystins were measured using UPLC-MS/MS using a Thermo Vanquish^TM^ UHPLC (Thermo Fisher Scientific, Waltham, MA, USA) quaternary pump system coupled to a Thermo Vanquish^TM^ Sample Hub and a Thermo TSQ Altis^TM^ Triple Quadrupole Mass Spectrometer to detect 11 common MCs (RR, YR, HtyR, [D-Asp^3^]-LR, LR, HilR, WR, LA, LY, LW, and LF) and nodularin-R. An acidified (0.1% formic acid) 30-70% aqueous acetonitrile gradient at a flow rate of 0.2 mL min^-1^ was used with a 2.1 x 35 mm Ace-3 (3μm) C-18 pre-column in series with a 1.0 x 100 mm Ace-3 C-18 column (MacMod Analytical, Chadds Ford, PA). MCs were quantified using a modified version of EPA method 544 (EPA, 2015) that adds MC-HtyR, [D-Asp^3^]-LR, HilR, WR, and LW as well as including one quantitation ion and two confirmation ions for each congener. H-ESI with a capillary voltage of 3.5 kV, ion transfer tube temperature of 325°C, and vaporizer temperature of 350°C was used with a flow of 50 arbitrary units (Arb) of sheath gas, 10 Arb auxiliary gas, and 1 Arb sweep gas. Additional method details including quantitation ions, fragmentation products, collision energies and RF lens voltages for each analyte can be seen in Supplemental Table 2. MCs were identified using the confirmation ion ratios and retention times as determined using standards (Supplemental Table 2). Detection limits varied between congeners and ranged from 0.006–0.475 (RR and WR respectively) µg microcystins L^-1^.

*Measurements of photosynthetic efficiency.* Three mL of chemostat culture were collected daily and maximum quantum efficiency (*F_v_/F_m_*) was measured using a Pulse-Amplitude Fluorometry (Phyto-PAM II, Heinz Walz GmbH) following the protocol published in *protocols.io* (Stark et al., 2022).

*Lake Erie microcosm experiments.* Field experiments were conducted at Ohio State University’s *F.T. Stone Laboratory* in Put-in-Bay, Ohio in August 2026. Surface water was collected in 20 L carboys at Maumee Bay St. Park Beach (Oregon, OH; 41.686107°N, -83.77194°W) on 19 August 2026. Homogenized lake water containing natural communities was distributed into eight 2-L polycarbonate bottles which were randomly assigned to two sunlight treatments (30 or 60 min; n = 4 bottles per treatment).

Bottles were incubated overnight at 26° C in a large water-filled, temperature-controlled mesocosm tanks and sampled immediately before sunrise (~6:00 AM) on 20 August 2026. Following the initial sampling, bottles were wrapped in black plastic to prevent light exposure and returned to the water incubator. At approximately solar noon (1:00 PM EST), bottles were removed the mesocosm. Three bottles from each treatment were exposed to direct sunlight (~1800 µmol photons m^-2^ s^-1^) for either 30 or 60 minutes, while one bottle from each treatment remained covered as a dark control. At each time point a 250 mL aliquot was collected from each bottle for toxin analysis as described above. Temperature was monitored using a thermometer placed in a separate bottle containing water acclimated to the same temperature as the experimental bottles to ensure that sunlight exposure did not cause overheating.

*Data visualization and statistical analyses.* All statistical analyses were performed in GraphPad Prism (v.10.6.1). Differences in microcystin concentrations across diel timepoints in continuous cultures were evaluated using repeated-measures ANOVA. For comparisons between dark and light periods, measurements from the three consecutive days were averaged within each cyclostat to yield a single value for each sampling period, with each cyclostat treated as an independent sample unit (n = 3). Differences between sampling periods were assessed using a paired t-test. For the sunlight exposure experiment using natural communities, destructive sampling and volume limitations prevented us from doing repeated measurements of the same bottle overtime, meaning an ANOVA could not be used. Instead, the endpoint measurement from each bottle was paired with its corresponding 0 min control. Two separate t-tests were performed to compare concentrations between 0 and 30 min and 0 and 60 min of sunlight exposure (n = 3). The relationship between microcystin concentration and exposure duration was also evaluated using a simple linear regression.

For raw data from the continuous culture and field microcosm study, see Supplemental Tables 3 and 4, respectively.
